# Optimal Decoding of Neural Dynamics Occurs at Mesoscale Spatial and Temporal Resolutions

**DOI:** 10.1101/2023.09.18.558322

**Authors:** Toktam Samiei, Zhuowen Zou, Mohsen Imani, Erfan Nozari

## Abstract

**Introduction:** Understanding the neural code has been one of the central aims of neuroscience research for decades. Spikes are commonly referred to as the units of information transfer, but multi-unit activity (MUA) recordings are routinely analyzed in aggregate forms such as binned spike counts, peri-stimulus time histograms, firing rates, or population codes. Various forms of averaging also occur in the brain, from the spatial averaging of spikes within dendritic trees to their temporal averaging through synaptic dynamics. However, how these forms of averaging are related to each other or to the spatial and temporal units of information representation within the neural code has remained poorly understood.

**Materials and Methods:** In this work we developed NeuroPixelHD, a symbolic hyperdimensional model of MUA, and used it to decode the spatial location and identity of static images shown to *n* = 9 mice in the Allen Institute Visual Coding - NeuroPixels dataset from large-scale MUA recordings. We parametrically varied the spatial and temporal resolutions of the MUA data provided to the model, and compared its resulting decoding accuracy.

**Results:** For almost all subjects, we found 125ms temporal resolution to maximize decoding accuracy for both the spatial location of Gabor patches (81 classes for patches presented over a 9x9 grid) as well as the identity of natural images (118 classes corresponding to 118 images). The optimal spatial resolution was more heterogeneous among subjects, but was still found at either of two mesoscale levels in nearly all cases: the area level, where the spiking activity of neurons within each brain area are combined, and the population level, where the former are aggregated into two variables corresponding to fast spiking (putatively inhibitory) and regular spiking (putatively excitatory) neurons, respectively.

**Discussion:** Our findings corroborate existing empirical practices of spatiotemporal binning and averaging in MUA data analysis, and provide a rigorous computational framework for optimizing the level of such aggregations. Our findings can also synthesize these empirical practices with existing knowledge of the various sources of biological averaging in the brain into a new theory of neural information processing in which the *unit of information* varies dynamically based on neuronal signal and noise correlations across space and time.

## INTRODUCTION

Neural dynamics span across a wide range of spatiotemporal scales, from (sub)cellular to regional and from (sub)millisecond to circadian and higher (Buzsaki, 2006; Bressler and Menon, 2010; Breakspear, 2017). Arguably, the most common link between neural dynamics across different spatiotemporal scales is averaging. Macroscopic measurements such as EEG, MEG, and fMRI reflect spatially-averaged activities of millions of neuronal post-synaptic potentials (Buzsáki et al., 2012; Logothetis et al., 2001) which are themselves the result of pre-synaptic spatial averaging through dendritic trees (Cash and Yuste, 1999) and are linked to higher-frequency spiking activity through synaptic temporal averaging (Kandel et al., 2013). Averaging is also the theoretical foundation for the broad family of mean-field models (Breakspear, 2017; Buice and Cowan, 2009), and is further applied across imaging modalities as a signal-processing step *for improving signal to noise ratio (SNR)* (Poldrack et al., 2011; Luck, 2014; Widmann et al., 2015). Averaging or averaging-involved methods such as spatial smoothing and parcellation of voxel-wise fMRI, low-pass filtering, principal component analysis (PCA), independent component analyses (ICA), peri-stimulus time histograms (PSTH), and firing rate estimations are all popular means for reducing the dimensionality of data and making large-scale brain recordings understandable and explainable.

On the other hand, averaging also involves an inevitable loss of information. This can be seen, at a generic level, from the information-theoretic data processing inequality (Cover, 1999). In a series of recent works (Nozari et al., 2023; Ahmed and Nozari, 2022, 2023), we have further shown that averaging has a particularly strong linearizing effect, transforming functionally-relevant nonlinearities (spiking, multi-stability, limit cycles, etc.) into what appears to be “noise” in macroscopic measurements. Notably, *the strength of this linearizing effect is directly related to the amount of signal correlation among the averaged units*: the higher signal correlation is among a group of neurons and the slower it decays with distance between them, the weaker the linearizing effect of averaging becomes, i.e., the more neurons we need to average over before nonlinearities fade (Nozari et al., 2023).

As such, averaging can have a dual effect on the neural code: it can improve SNR by averaging over noise, but it can also degrade SNR by cancelling out functionally-relevant nonlinearities. The balance of these two effects depends on the relative strength of signal and noise correlations among neurons. *If* noise correlations are weaker and decay more rapidly with distance, then controlled amounts of averaging can be beneficial by cancelling noise faster than fading the signal. Otherwise, no amount of averaging would be beneficial and the neural code can be best decoded from the raw spiking activity of individual neurons with millisecond resolution.

In this work, we test the *central hypothesis* that there exists an optimal amount of spatial and temporal averaging, i.e., an optimal spatiotemporal resolution, which maximizes neuronal SNR and therefore the accuracy of decoding the neural code. Using data from *n* = 9 mice from the Allen Institute Visual Coding - Neuropixels dataset, we design computational models that classify visual images shown to each mouse using its large-scale MUA with parametrically varied amounts of spatial and temporal averaging. The use of the brain-inspired hyper-dimensional computing (HDC) framework (Kanerva, 2009; Schlegel et al., 2022; Zou et al., 2022) allows us to gain precise control over the amount of end-to-end spatiotemporal averaging performed by the decoder and minimize implicit sources of averaging that extensively occur during the training of most machine learning alternatives and can confound our findings. The resulting HDC-based classifier, termed NeuroPixelHD, provides a means to testing this work’s central hypothesis as well as a general-purpose model for encoding and decoding large-scale MUA data in a transparent and interpretable manner owing to the symbolic nature of HDC.

## RESULTS

### NeuroPixelHD: a Hyperdimensional Model for Large-Scale Multi-Unit Activity

In this work we use the brain-inspired framework of HDC (Kanerva, 2009; Schlegel et al., 2022; Zou et al., 2022) to design NeuroPixelHD, an efficient decoding model for MUA. The use of HDC to test our central hypothesis is motivated by the core observation that vector summation results in an *irreversible averaging* in small dimensions (i.e., the summands are not recoverable from the sum), but it can result in *reversible memorization* in very large dimensions (Supplementary Note 1). As such, a trained HDC model can embed a copy of all of its training samples, without any unintended implicit averaging. In this work, we train NeuroPixelHD to classify images within two categories based on MUA recordings: Gabor patches at different locations of a 9x9 grid in the visual field, and 118 different images of natural scenes (Methods).

Inspired by our earlier work on event-based cameras (Zou et al., 2022), the design of NeuroPixelHD involves an encoding phase and an adaptive training phase. During the encoding phase, the binned spike counts of all the recorded neurons throughout each trial (250ms here) is encoded into one hyper-vector (HV) (Figure 1). As described in details in Methods, The encoding involves a sequence of reversible binding and bundling operations (standard in HDC, see Methods) over three ingredients: binned spike counts, neuron HVs, and time bin HVs. Neuron HVs are generated based on each neuron’s anatomical region and response during receptive field tuning, therefore maintaining a level of spatial correlation proportional to the anatomical and functional proximity of each pair of neurons. Time bin HVs are generated randomly for the beginning and end of each trial (0ms and 250ms) and interpolated via linear dimension borrowing for intermediate bins, maintaining a level of temporal correlation proportional to the temporal proximity of each pair of time bins. These HVs are then fused with binned spike counts using binding and bundling operations to encode all the neural activity during each trial into one trial activity HV used during the second phase for classifier training.

**Figure 1.**
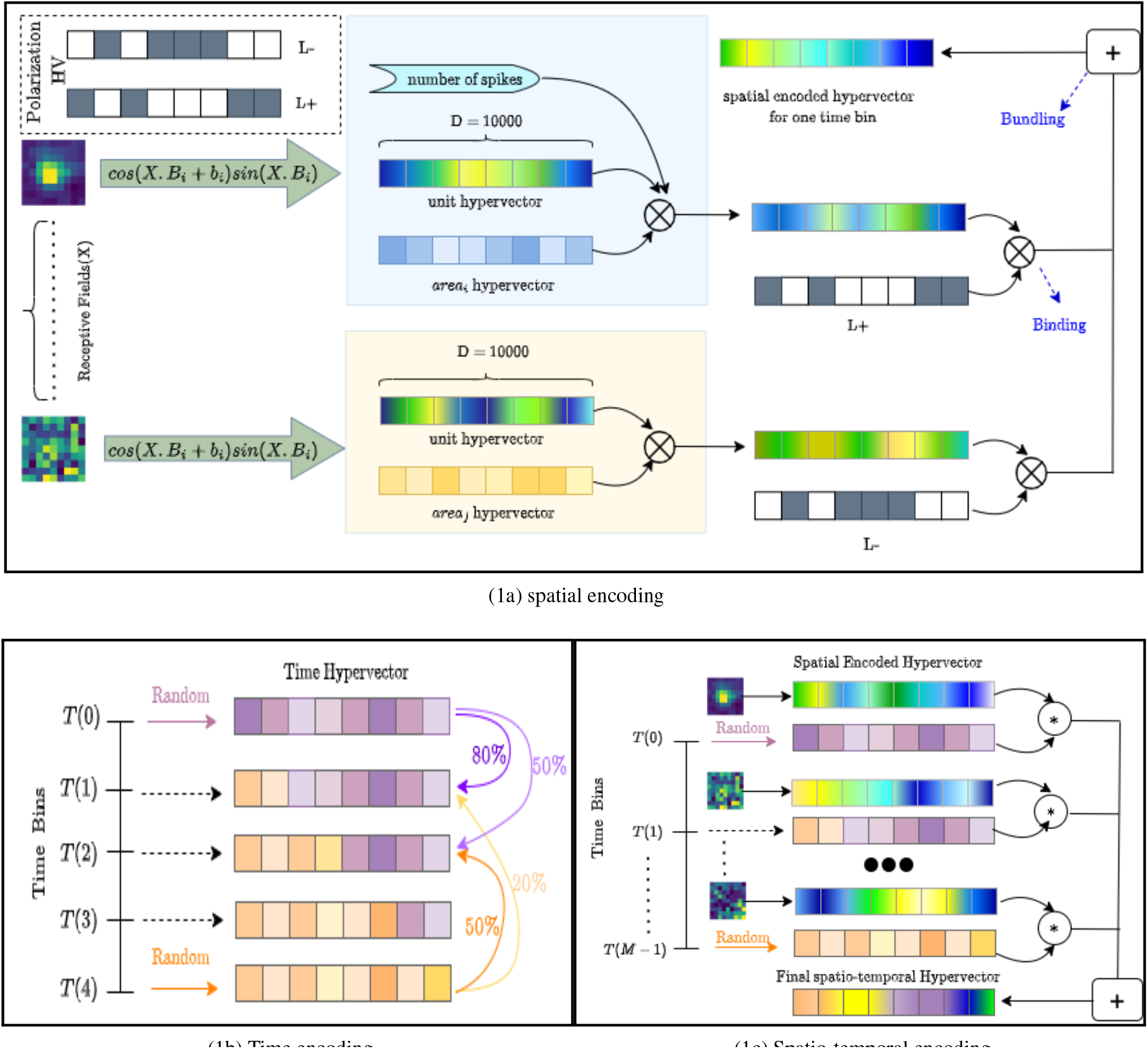
The structure and encoding of NeuroPixelHD. **(1a)** Spatial encoding in NeuroPixelHD. Spatially correlated hypervectors are generated for each neuron, using each neuron’s responses to receptive field tuning and cosine encoding, and then bound with (randomly generated) hypervectors representing corresponding brain areas. **(1b)** Temporal encoding in NeuroPixelHD. Two random hypervectors are generated for times 0 and 250ms and then linearly interpolated, via dimension borrowing, to generate correlated hypervectors for intermediate time points. **(1c)** Encoding of each trial in NeuroPixelHD.

The second phase of NeuroPixelHD consists of adaptive training. Each class (Gabor location or natural scene image) is represented by one class HV. All class HVs are initialized at zero and iteratively updated such that the similarities between each class HV and corresponding trial activity HVs gradually increase and the similarities between each class HV and trial activity HVs of other classes gradually decrease (see Methods). We use cosine similarity (normalized dot product) in NeuroPixelHD due to its simplicity and computational efficiency, but various other measures of similarity have also been proposed in HDC and can be alternatively used. At the end of training, each test trial is assigned to the class that has the largest similarity between its class HV and activity HV of that test trial. To measure classification accuracy, we use standard F1 score for natural scene images and median Euclidean error between the actual and predicted locations for Gabor patches (see Methods).

We compared NeuroPixelHD against state-of-the-art classification models to ensure that our focus on minimizing its implicit sources of averaging has not compromised its classification performance. In comparison to random forests, support vector machines, and artificial neural networks, NeuroPixelHD achieved comparable classification accuracy and efficiency (Figure 2a). This comparison becomes more favorable for NeuroPixelHD at finer temporal resolutions, where the input features for classification become larger dimensional. At the finest resolution (largest feature dimensions), NeuroPixelHD is notably more efficient than other methods while achieving comparable or better accuracy (Figure 2b,2c). These result confirm the utility of NeuroPixelHD as an accurate and efficient classification model for MUA. In what follows, we will then use NeuroPixelHD to probe into the costs and benefits of averaging and the optimal spatiotemporal resolution for decoding information from MUA.

**Figure 2.**
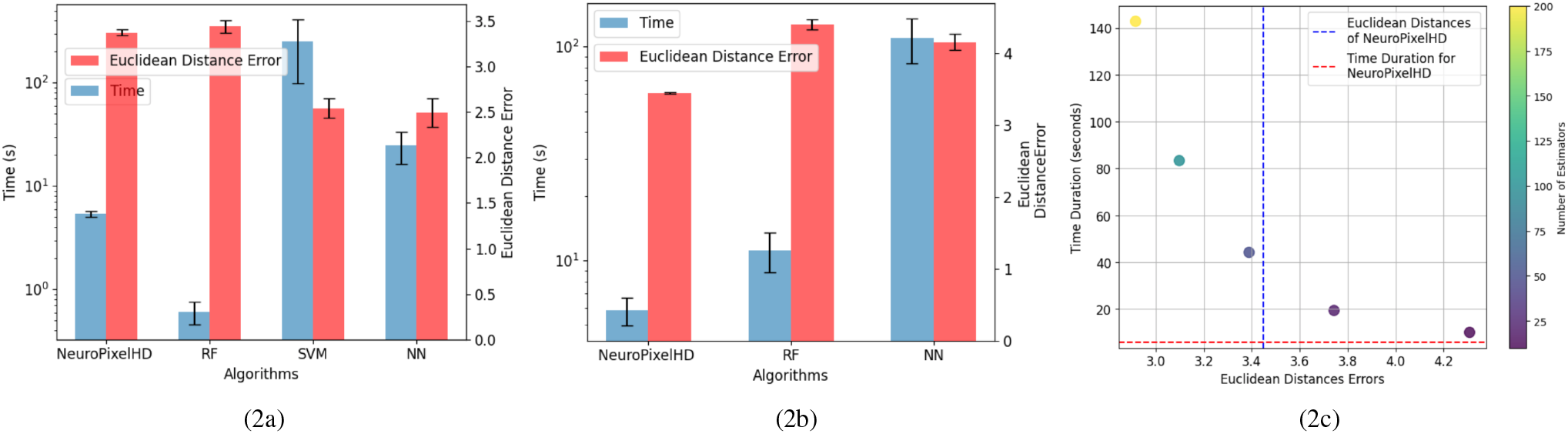
NeuroPixelHD has comparable performance with standard machine learning classifiers. All models are trained for classifying the location of Gabor stimuli (cf. Methods), separately for *n* = 9 mice. **(2a)** Mean time complexity (CPU time) and Euclidean distance error of NeuroPixelHD, support vector machine (SVM), random forest (RF), and neural networks. Note the logarithmic scaling on the left vertical axis. All models are compared using a 10ms time resolution at which the dimension of the input vectors to different models is very similar. **(2b)** Same as (2a) but using a 1 ms time resolution. The run time for SVM was significantly longer than other methods (over 8 hours) and we thus excluded SVM from this comparison. **(2c)** Scatter plot illustrating a more detailed relationship between time complexity and Euclidean distance error of NeuroPixelHD and RF with varying number of estimators (trees) within a 1 ms time resolution. While RF can achieve higher accuracy than NeuroPixelHD with sufficiently large number of estimators, NeuroPixelHD achieves a better complexity-accuracy trade off.

### 125ms temporal resolution maximizes visual decoding accuracy for static images

We investigated the optimal amount of temporal averaging for visual decoding by comparing the decoding accuracy of NeuroPixelHD for varying bin size values. We started from the smallest bin size of 1ms and gradually increased the bin size until reaching one bin for the entire trial duration (250ms). As we increase the bin size, both the signal and the noise components of spike counts are averaged, potentially changing the spike counts’ signal to noise ratio and, in turn, the decoding accuracy of the downstream classification.

When classifying the location of Gabor patches from binned MUA spike counts, in most subjects, we observe an initial insensitivity of classification accuracy to bin size between 1-10ms, followed by a sharp improvement in decoding accuracy until 125ms, and an occasional worsening of accuracy afterwards (Figure 3a). Remarkably, for most subjects, the worst accuracy occurs at the smallest bin size, despite the classifiers’ access to *all* spike count information. This may be at first counter-intuitive from an information-theoretic perspective (cf., e.g., the Data Processing Inequality (Cover, 1999)), but demonstrates the importance of optimal feature extraction from a machine learning perspective and is consistent with the common perception of individual spikes as being highly noisy and the common practice of binning spike counts before using them for downstream analyses.

**Figure 3.**
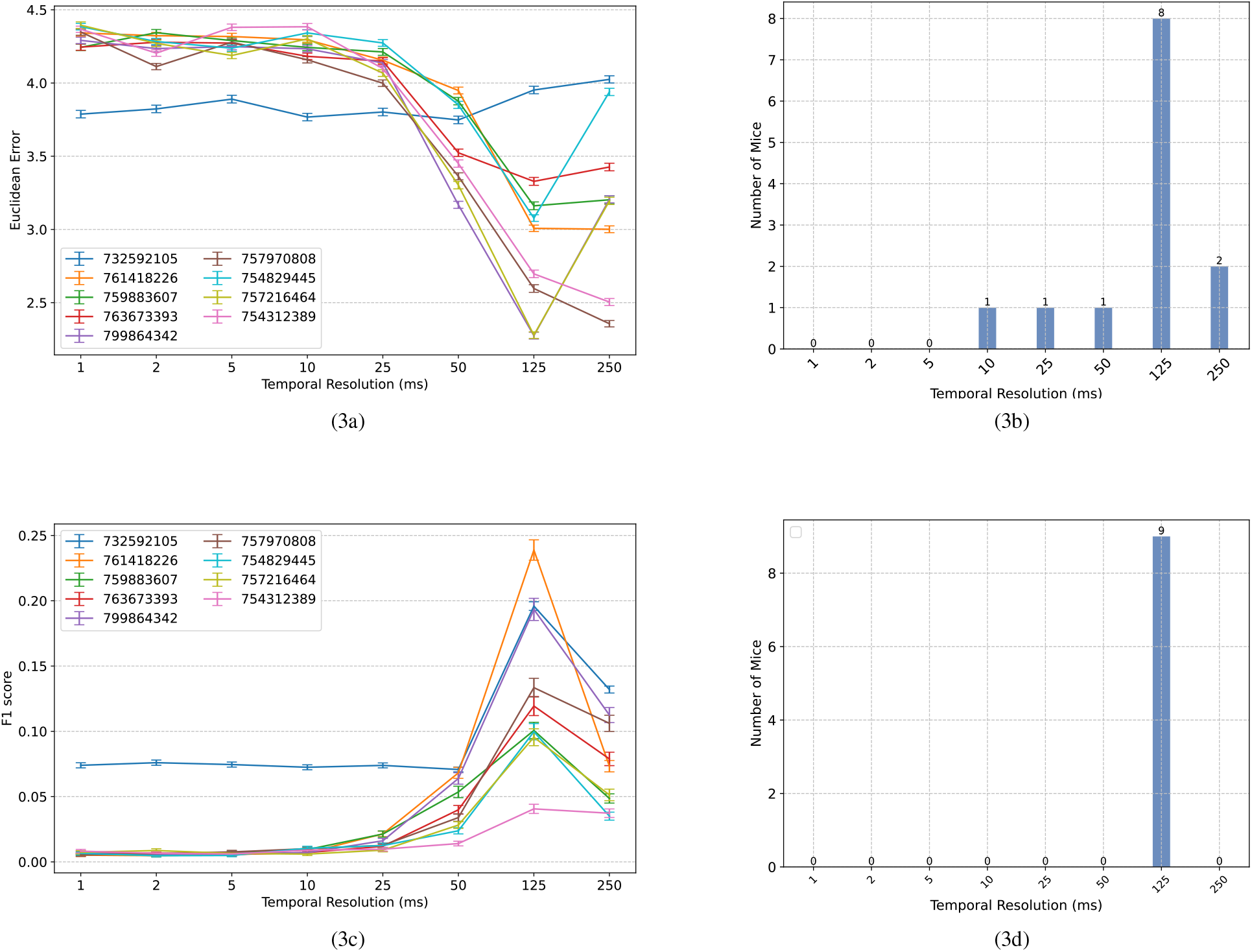
Comparisons between classification accuracy of NeuroPixelHD with different temporal resolutions. **(3a)** Mean Euclidean distance errors of NeuroPixelHD in classifying Gabor locations. Each line corresponds to one mouse (*n* = 9) and error bars represent 1 s.e.m. **(3b)** Distribution showing the number of mice for whom each time bin is optimal. The optimal time bin for each mouse was selected based on Wilcoxon signed rank test with *α* = 0.05. In cases where 2 or more bin sizes had the least error (insignificant statistical difference), all of them were counted as optimal bin size for that mouse and included in the aggregate bar graph. **(3c**,**3d)** Similar to (3a,3b) but for the classification of nature scenes. Here accuracy is measured by F1 score (higher is better; see Methods). Across both tasks, the 125ms time bin resulted in maximum decoding accuracy.

To further resolve the heterogeneity among subjects and compute the optimal temporal resolution at the group level, we found the optimal bin size for each subject (namely, the bin size with the lowest median classification error) and calculated, for each bin size, the number of subjects for whom that bin size is optimal. If two or more bin sizes were jointly optimal (*p* ≥ 0.05, Wilcoxon signed rank test), we included all of them in the group-level count. The result, shown in Figure 3b, corroborates that 125ms resolution is optimal at the group level, 250ms is the second best, and 1-5ms resolution yields the least signal to noise ratio overall. The same trend appears even more contrastively for the decoding of natural scenes (Figure 3c,3d). Here, we measure classification accuracy using F1 score with higher values indicating higher accuracy. Across all subjects, the 125ms resolution provides the highest decoding accuracy, while the 1-10ms resolutions result in chance level classification (1/118 ≃ 0.008) in all but one mouse.

### Population and Area Level Spatial Resolutions Maximize Visual Decoding Accuracy

We next performed a dual analysis, comparing the visual decoding accuracy of NeuroPixelHD when the spike counts provided at its input were spatially averaged at progressively larger scales. We used five levels to divide the range from micro to macro scale: single neuron level, where no averaging is performed; population level, where the spike counts of regular spiking (putatively excitatory) and fast spiking (putatively inhibitory) neurons within each brain area were combined; area level, where the spike counts of all neurons within each area were combined; region level, where the spike counts of all neurons within all areas of each brain region were combined; and whole-brain level, where the spike counts of all recorded neurons were combined (cf. Methods). The classification accuracy of NeuroPixelHD was then compared between these levels for each mouse, separately for the Gabor patches and natural scenes.

Similar to the above analysis of temporal averaging, the optimal resolution was at the micro nor at the macro scales, but rather at an intermediate (meso) scale. For the classification of the spatial location of Gabor patches, for almost all subjects, maximum decoding accuracy (minimum Euclidean error) was obtained at either the population level or the area level (Figure 4a). In particular, the two extremes of neuron and whole-brain levels are significantly worse than the intermediate levels and not optimal in any of the subjects (Figure 4b). The same trend also appeared in the decoding of images of natural scenes. For all but one subject, NeuroPixel’s classification accuracy (measured via F1 score, see Methods) was at the chance level 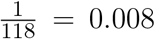 at the neuron and whole-brain levels and reached its maximum at an intermediate level (Figure 4c). In fact, in most subject, the maximum decoding accuracy was obtained at either the population or the area level as was the case in the Gabor task (Figure 4d).

**Figure 4.**
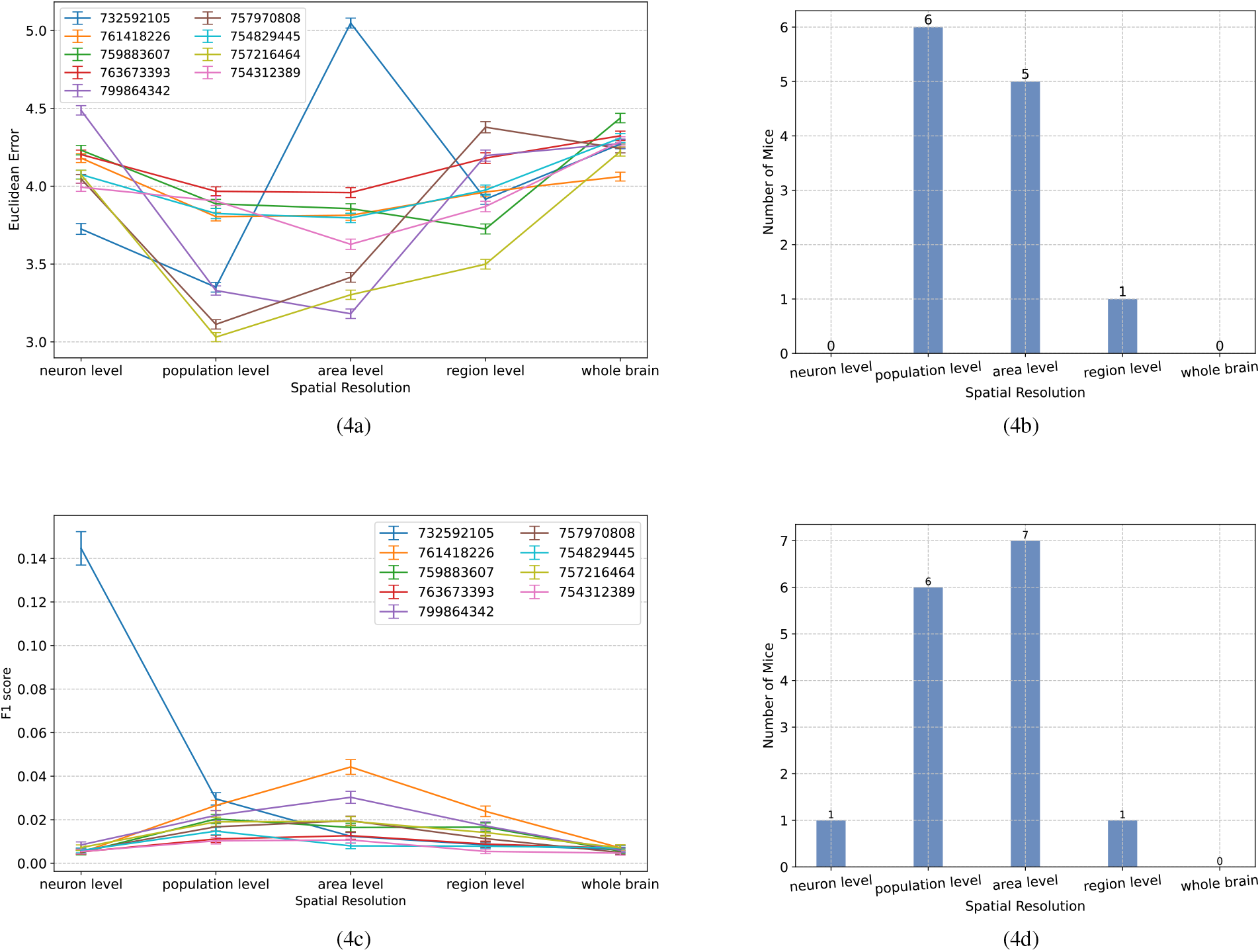
Comparisons between classification accuracy of NeuroPixelHD with different spatial resolutions. **(4a-4d)** Panels parallel those in Figure 3 except that averaging is performed over space (clusters of neurons). See Methods for a description of each spatial scale. All comparisons are performed without temporal averaging (1ms time bin). Across both tasks and most subject, either the population level or the area level resolutions led to maximum visual decoding accuracy.

In summary, across both the temporal and spatial dimensions of visual coding as well as spatial localization and object identification, we found intermediate resolutions, rather than the micro or macro extremes thereof, to maximize decoding accuracy in most cases. This is consistent with a model in which noise correlations decay more rapidly among nearby neurons than do signal correlations, and confirms our initial hypothesis that averaging initially improves, but then degrades, neuronal SNR and therefore the accuracy of decoding the neural code.

## 1 DISCUSSION

In this study we designed NeuroPixelHD, a normative hyperdimensional computational model for large-scale MUA, and used it to probed into the effects of spatial and temporal averaging on the neuronal signal to noise ratio in the brain. While the largely averaging-free architecture of NeuroPixelHD was its key property in allowing us to gain precise control over the amount of end-to-end averaging performed by the model, we also demonstrated its comparable accuracy and higher efficiency compared to standard machine learning models in large dimensions. We compared the decoding accuracy of NeuroPixelHD when its input spike counts were averaged to varying degrees over space and time, and found 125ms temporal resolution and population-area level spatial resolution to maximize the accuracy of decoding both the spatial location of Gabor patches and the identity of natural scenes from large-scale MUA.

An interesting finding of this study was a confirmation of the widespread belief that neural populations clustered based on cell type form functionally relevant units for studying the neural code (Klausberger and Somogyi, 2008; Pfeffer et al., 2013; Jadi and Sejnowski, 2014). However, our results also show that in the absence of ground-truth genetic information, this is a nuanced clustering sensitive to the functional proxy used for cell type differentiation. Putatively excitatory and inhibitory neurons are often interchangeably classified based on their spiking waveform shape or spiking statistics (Tseng and Han, 2021; Becchetti et al., 2012; Connors and Gutnick, 1990; Barthó et al., 2004). However, we found the two proxies to lead to notably distinct clusters (Supplementary Figure 1b) and clustering based on Fano factor to give significantly better classification results (Supplementary Figure 1c,1d). This marked difference is in need of further mechanistic investigation, but in itself highlights the importance of functional proxies used for population-level analysis of neural dynamics.

An unconventional aspect of NeuroPixelHD encoding is the use of independent time bin HVs for different trials (even though time bin HVs within each trial are correlated, cf. Methods). This is critical for preventing averaging to occur among trial HVs during the adaptive training process where HVs of different trials are linearly combined (bundled). Using shared time bin HVs would instead result in every pair of trial HVs to become more similar to each other, due to the shared spatial and magnitude similarity within the same time bin. This is the similarity preserving property of binding: *δ* (*a ⊗c, b ⊗c*) = *δ* (*a, b*). In comparison, When independent time bin HVs are applied, the same similarity no longer transfers to similarity between trial HVs, leading to a broader and more widespread usage of the hyperspace (cf. Supplementary Note 1). On the other hand, using *shared* time HVs leads to an improved classification accuracy (Supplementary Figure S2). This is expected, particularly in light of the benefits of moderate amounts of averaging that we observed (cf. Results), but is still undesirable for the purposes of this study as the implicit averaging implied by using shared time bin HVs can confound the explicit amounts of averaging we perform at each spatiotemporal resolution and potentially bias our results.

Similarly, it is important to disentangle the strengths and weaknesses of NeuroPixelHD compared to other machine learning models (RF, SVM, etc.). In general, HDC-based models may at times achieve higher task accuracy compared to non-symbolic alternatives (Kim et al., 2018; Imani et al., 2017), but their main strength lies in their symbolic and interpretable structure and computational efficiency (Kleyko et al., 2023; Thomas et al., 2021; Imani et al., 2021). The interpretability of HDC allows one to ask “cognitive” queries from a learned model. This includes (1) memory recall, as in decoding spiking activities from the HV, (2) pattern recognition, as in recognizing certain spatial-temporal patterns of spiking activities, and (3) feature attribution, where one can backtrack which set of spiking activities in a query HV produces the highest similarity signal to the class HVs, an analytical approach commonly applied for explainable machine learning (Lundberg et al., 2018) with the additional benefit that the HDC representation of spatial and temporal features are explicit. The key trick to leveraging HDC cognitive computation is to realize that every transformation by an HDC operator can be interpreted as a certain type of data structure manipulation - (weighted) bundling creates (weighted) sets and binding creates association (Kanerva, 2009) - and that the similarity between two HVs reflects their structural alignment. HDC models are also commonly more efficient than classical alternatives, making them able to handle larger data sets and/or converge in fewer epochs (Ge and Parhi, 2020) (cf. Figure 2b,2c). The fact that HDC operations (bundling, binding, and permutation) can be averaging-free and reversible further adds to its advantages for the purposes of this study. Finally, we should emphasize that NeuroPixelHD is the first HDC-based classifier designed for NeuroPixel data, and thus may not be the best one. Future studies are needed to investigate the full potential of HDC in encoding and decoding large-scale MUA data.

This study has also some limitations. As noted earlier, NeuroPixelHD is not necessarily the best HDC architecture for encoding and decoding large-scale MUA data. Our analysis is further limited to two categories of visual stimuli, making it possible that other, possibly very different, spatial and temporal resolutions are optimal for different categories of stimuli, modalities, and tasks. Moreover, our analyses of optimal spatial resolution is likely confounded by the sparse sampling of neurons in our dataset. Should we had access to spiking activity of all neurons in each region, we might have found different, possibly finer, resolutions to be optimal for decoding.

Overall, this study presents empirical support for the presence of an optimal amount of spatial and temporal averaging that maximizes the neuronal signal to noise ratio, and provides an initial estimate of optimal spatial and temporal resolutions during passive viewing of static images. Future work is needed to extend these estimates to other tasks, sensory modalities, and species. Further investigations are also necessary to relate the data-driven estimates found in this study to the underlying biological mechanisms, including synaptic time constants, axonal conduction velocities, and signal and noise correlations among populations of excitatory and inhibitory neurons. Finally, it remains an invaluable area of future research to understand the relationship between the spatiotemporal resolutions that are optimal for a normative decoding model such as NeuroPixelHD and those that are optimal for and/or employed by the brain itself.

## MATERIALS AND METHODS

### Visual Coding – Neuropixels Dataset

In this study, we utilized data from the Allen Brain Observatory, specifically from experiments conducted with Neuropixel probes in wild-type mice. The initial Neuropixels data release encompassed responses from neurons in the visual cortex, hippocampus, and thalamus, including brain regions such as: Striate Cortex, Dorsal Extrastriate Cortex, Ventral Extrastriate Cortex, Hippocampus, Subiculum, Dentate Gyrus, Thalamus, Hypothalamus, and Midbrain.

Different visual stimulation tasks were administered to mice, as illustrated in Figure 4. However, for our data analysis, we focused on two specific tasks: Gabor and natural scenes. All experimental sessions commenced with a receptive field mapping stimulus. During the Gabor task, Gabor patches were randomly displayed at one of 81 locations on the screen, forming a 9 x 9 grid. Each patch appeared for 250 ms, without any blank intervals, and this process was repeated 45 times for each location.

For the natural scenes task, a stimulus comprising 118 grayscale natural images was employed. These images were sourced from the Berkeley Segmentation Dataset (Martin et al., 2001), the van Hateren Natural Image Dataset (van Hateren and van der Schaaf, 1998), and the McGill Calibrated Colour Image Database (Olmos and Kingdom, 2004). Prior to presentation, the images underwent contrast normalization and resizing to 1174 x 918 pixels. Each image was randomly shown for 0.25 seconds, without any intervening gray period. For this task, each image was shown 50 times.

### HyperDimensional Computing (HDC)

In HDC, “hypervectors” (HVs), i.e., high-dimensional representations of data created from raw signals using an encoding procedure, constitute the basic building blocks of computational algorithms (Kanerva, 2009). These hypervectors are then combined and manipulated using specific mathematical operations (see below) to build transparent, symbolic computational models with the ability to preserve (memorize) the original information. Such memorization is enabled by a key property called “near-orthogonality”. Consider two HVs 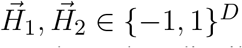 whose elements are independent and identically distributed (i.i.d.), each following the Rademacher distribution. If *D* is large enough (often *D* ∼ 10^4^ in practice), these vectors become approximately orthogonal, as can be seen from their cosine similarity

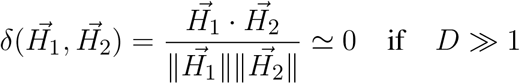

As such, (pseudo)random HVs with i.i.d. components are commonly used as essential ingredients in HDC encoding processes. Such HVs are then combined using established HDC operations to generate new HVs that have compositional characteristics and therefore allow computations to be performed in superposition, effectively encode spatial and temporal information, and respect intricate hierarchical relationships present in the data. The most commonly used HDC operations in the literature are as follows (Gayler, 1998; Zou et al., 2022; Kleyko et al., 2023):

### Binding (⊗)

Two HVs are bound together using component-wise multiplication of their elements. This operation is often used for creating association among HVs, is reversible 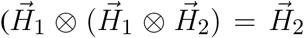 and 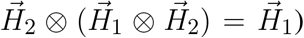, and the resulting HV can be shown to be nearly orthogonal to both operands 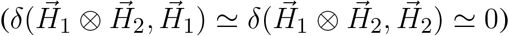.

### Bundling (+)

Two HVs are bundled together using component-wise addition of their elements. Unlike summation in small dimensions which results in an (irreversible) averaging, hyperdimensional bundling preserves the information of both operands. This can be seen from the fact that the bundled HV has non-negligible similarity with each of its operands 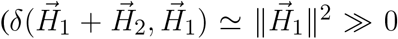 and 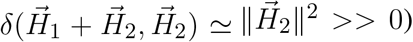. Therefore, by performing a similarity check between a bundled HV and any query HV, one can determine whether the query has been one of the constituents of the bundled HV.

### Permutation (*ρ*)

Permutation is achieved by a circular shift of one HV’s elements and is used to generate sequential order among HVs. We do not use permutation in the encoding of NeuroPixelHD.

### NeuroPixelHD Encoding

#### Receptive field encoding

In this study, we adopted a novel approach to encode the identity of each neuron into one HV. Unlike using i.i.d. HVs for different neurons, this approach generates neuron HVs which are correlated with each other depending on the similarity between the receptive fields of their corresponding neurons. For this, we used the Gabor receptive field tuning experiments and computed the mean spike count of each neuron during the full 250ms presentation of each of the 81 Gabor locations, averaged over the 45 repetitions of each location. This generates a (pre-encoding) 81-dimensional receptive field response vector 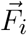 for each neuron *i*, which is then encoded into a *D*-dimensional HV via (Hernández-Cano et al., 2021; Rahimi and Recht, 2007)

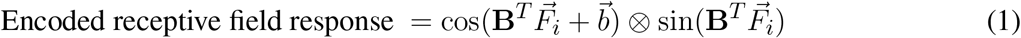

where **B** is a 81-by-*D* random matrix with i.i.d. standard normal elements, 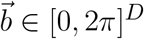 is a random vector with i.i.d. elements uniformly distributed over [0, 2π], and *D* = 10^4^ ≫ 81. This encoding is inspired by the Radial Basis Function (RBF) kernel trick and can account for nonlinear relationships among features during encoding.

#### Brain area encoding

In each of the subjects, spiking data from neurons in a subset of the following brain areas was available: VISp, VISam, VISal, VISrl, VISmma,VISpm, VISl,CA1, CA2, CA3,SUB, ProS,DG,TH, LP, LGv, LGd, PP, PIL, MGv, PO, Eth, POL, ZI, and APN. In principle, neurons in distinct areas can have very similar receptive field responses. Therefore, to further distinguish neurons from different areas, we define a unique, random and independent HV 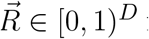 for each brain area. To simplify indexing notation, we use 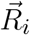 to denote the area HV corresponding to each neuron *i*, thus 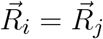 if neurons *i* and *j* belong to the same area. These area-specific HVs are then bound with encoded receptive field HVs, as described below (cf. Equation (2))

#### Spiking activity encoding

In each time bin, each neuron may have a zero or non-zero number of spikes. Both the occurrence and absence of spikes contains valuable information which need to be reflected in overall trial encoding. Motivated by our prior work (Zou et al., 2022), we define two *polarization* HVs 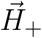 and 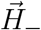, corresponding to the presence and lack of spikes, respectively. We generated 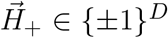 with i.i.d. elements and let 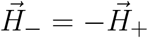.

#### Time encoding

The duration of each trial, set at 250 milliseconds, is divided into *B* bins, *B* = 1, 2, 5, 10, 25, 50, 125, 250. For each time bin,, *t* = 0, 1,…, *M* 1, which *M* is the number of time hypervectors 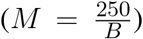, a 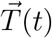 is constructed such that temporal correlation is maintained among 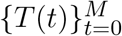. This is achieved, independently for each trial, by generating random 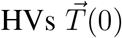 and 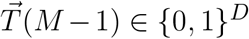 for the initial and final time bins and linearly interpolating between them to generate time HVs for intermediate bins. Mathematically,

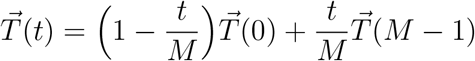

The resulting HVs retain temporal relationships depending on their temporal proximity.

#### Trial encoding

Finally, the HVs described earlier are combined through various levels of binding and bundling to generate a single HV encoding of each trial. This is done via

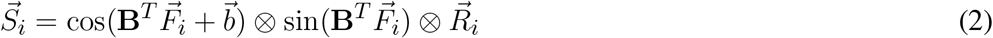

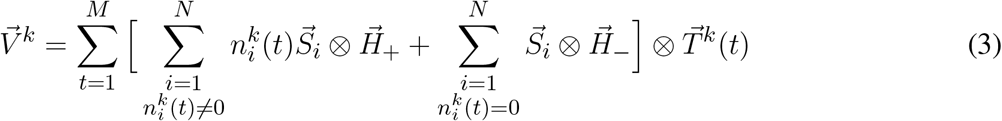

The spatial HV 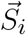 is the encoding (i.e., identity) of each neuron *i* and results from binding its encoded receptive field response in Equation (1) with its encoded area HV 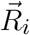. These spatial HVs are then scaled and polarized appropriately, bundled over space, bound with corresponding time HVs, and then bundled over time to generate the final trial HV 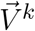.

### NeuroPixelHD Adaptive Training

Following our earlier work (Zou et al., 2022), we employed an adaptive training approach that considers the *extent* to which each training data point is correctly or incorrectly classified in updating the class HVs. Consider a problem with *m* classes and *K*_train_ training samples (represented by encoded trial HVs) 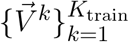, where each training sample *k* ∈ 𝒞 _*l*_ for some class *l* = 1,…, *m*. 𝒞 _*l*_ denotes the set of all trial indices that belong to class *l*. The goal of the training is to generate one class HV 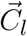 for each class *l* = 1,…, *m* such that each test sample 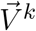 has the highest similarity with its own class HV, i.e.,

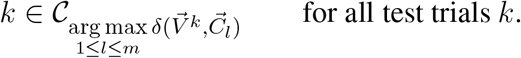

All class HVs are initialized to zero at the beginning of training and gradually updated such that their similarity with training samples of their own class is increased and their similarity with training samples of other classes is decreased.

Consider first the case for the classification of natural scenes (*m* = 118). At initialization, all 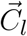 are set to zero. Then, for each training sample 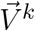, let ℓ denote its correct class (*k* ∈ 𝒞 _*ℓ*_) and ℓ′ denote its predicted class 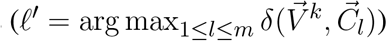. Further, define

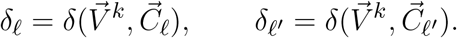

When the training sample is predicted correctly (ℓ′ = ℓ), the correct class HV 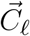 is updated in order to further increase its similarity with 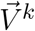:

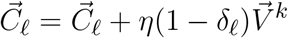

The update is proportional to 1 − δ_*ℓ*_ so that 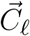 is modified less if its similarity with 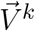 is already high. If the training sample is predicted incorrectly (ℓ′ ≠ ℓ), the predicted class is also updated such that its similarity with 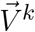 is decreased,

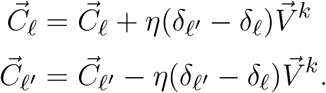

Similar to the previous case, the adaptive training considers the extent to which a training point is misclassified. In cases where the prediction is significantly off (δ_*ℓ*′_ ≫ δ_*ℓ*_) the update equation substantially modifies 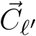, whereas for marginal mispredictions (δ_*ℓ*’_ ≃ δ_*ℓ*_), the update makes smaller adjustments. For both cases, we used *η* = 0.01 and performed the training for 3 epochs (rounds of presenting the training samples).

The above equations are slightly adjusted for the classification of the location of Gabor patches (*m* = 81) due to the presence of a natural notion of proximity between classes. In this case when the query data is predicted correctly (ℓ = ℓ′), we update not only the correct class HV 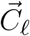 but also the HVs for the (up to 8) classes adjacent to it, i.e.,

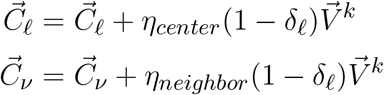

for all classes (Gabor locations) *ν* adjacent to ℓ. Similarly, when each training sample is predicted incorrectly (ℓ′ ≠ ℓ), we let

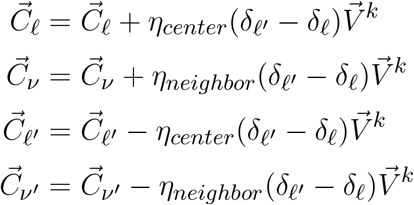

for all classes *ν* adjacent to ℓ and *ν*′ adjacent to ℓ^*’*^. We used *η*_*center*_ = 0.01 and *η*_*neighbor*_ = 0.001 and executed the algorithm for 2 epochs.

### Spatial Averaging

In this study, we employed the HDC algorithm at various levels of spatiotemporal resolution. The spatial averaging involved five, progressively coarser levels: neuron level, population level, area level, region level, and whole brain. At the neuron level (no spatial averaging), neurons were the basic spatial units, and we used the spike counts of all recorded neurons separately during the trial encoding process as summarized in Equation (2) and Equation (3).

At the population level, we clustered the neurons within each brain area into putatively excitatory (E) and inhibitory (I) populations. Among the various functional proxies suggested for E/I classification (Connors and Gutnick, 1990; Barthó et al., 2004; Becchetti et al., 2012), we used spike count Fano factor found in (Becchetti et al., 2012) to nearly perfectly distinguish the two populations. For each neuron, its Fano factor (variance-to-mean ratio) was computed based on its number of spikes during a specific time bin across all Gabor positions and trials. This value is expected to be higher for inhibitory neurons than excitatory ones (Becchetti et al., 2012). Therefore, given the nominal 80-20 ratio of excitatory and inhibitory neurons (Markram et al., 2004; Beaulieu, 1993), we labeled the 20% of neurons in each area with highest Fano factor as putatively inhibitory and the rest as putatively excitatory (Supplementary Figure 1a). The spike counts of neurons within each E/I population were than summed and used instead of 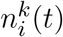 in Equation (3), where *i* now refers to a population rather than a neuron. Accordingly, the spatial HVs 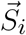 in Equation (2) were also replaced by population HVs computed via binding a randomly generated E/I HV (same across all regions) with the corresponding area HV 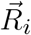. Considering the potential for the above clustering based on Fano factor to produce varied clusters depending on the chosen time resolution, particularly for neurons that exhibit intermediate traits, we compared the classification accuracy of population-level classifiers that used different bin sizes in the computation of Fano factors, and selected the Fano factor bin size that achieved the highest classification accuracy for each mouse. The resulting optimal population-level model was then compared against other spatial resolutions at the finest (1ms) temporal resolution.

At the area level, the spike counts of all neurons within each area were summed and used instead of 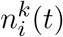 in Equation (3). Spatial HVs 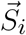 in Equation (3) were accordingly replaced by 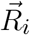. Similarly, at the region level, the spike counts of all neurons within each region were summed and used instead of 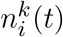 in Equation (3). Table 1 shows the assignment (clustering) of brain areas to regions. For each region, one random and independent spatial HV was generated and used as 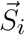 in Equation (3). Finally, at the whole-brain level, the spike counts of all recorded neurons for each mouse were combined, simplifying Equation (2) and Equation (3) to 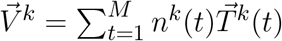.

**Table 1.** List of brain areas available within each brain region. Each mouse has data from neurons in some subset of these areas. The “dorsal” and “ventral” extrastriate cortices are named as such for ease of reference and based on homology to the primate brain (Marshel et al., 2011), not anatomical location in the mouse brain (cf. (Institute, 2019, Fig. 6)).

| Region | Included areas |
| --- | --- |
| Striate Cortex | VISp |
| Dorsal Extrastriate Cortex | VISam, VISal, VISrl, VISmma |
| Ventral Extrastriate Cortex | VISpm, VISl |
| Hippocampus | CA1, CA2, CA3 |
| Subiculum | SUB, ProS |
| Dentate Gyrus | DG |
| Thalamus | TH, LP, LGv, LGd, PP, PIL, MGv, PO, Eth, POL |
| Hypothalamus | ZI |
| Midbrain | APN |

### Temporal Averaging

To assess the optimal temporal resolution for visual decoding, we binned raw spike counts into bin sizes of 1, 2, 5, 10, 25, 50, 125, 250ms, effectively averaging spike counts at the finest scale (1ms) over larger bins. The resulting binned spike counts are provided to the NeuroPixelHD encoder, i.e., 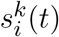 in Equation (3).

### Data Augmentation

The amount of neural data available to train the NeuroPixelHD classifier is relatively small compared to contemporary machine learning experiments, even though it consists of one of the largest MUA datasets available to date. In particular, the number of trials in which the exact same stimulus is shown to the mice (45 for Gabor patches and 50 for natural images) allows for no more than 1-2 dozen test samples per class, which often results in low statistical power when comparing among different spatiotemporal resolutions. This is often treated with data augmentation, for which various techniques have been proposed (Shorten and Khoshgoftaar, 2019; Antoniou et al., 2017; Bayer et al., 2022). In this work we used a novel form of data augmentation for comparisons between different spatial and temporal scales which exploits the specific dynamical structure of our data. Let the three-dimensional array **N**_*N ×T ×K*_ contain all the binned spike counts of *N* neurons over *T* time bins and *K* trials of the same class (same image). Then, we randomly shuffle the trial indices, uniformly for all neurons and independently for all times. In other words, we generate *T* independent sets of random indices 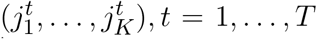 each of which is a permutation of (1,…, *K*), and generate a permuted array 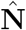 where

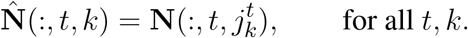

This process is repeated 4 times for each class, separately among training and test samples, resulting in a 5-fold increase in the total number of training and test samples.

### Measures of classification accuracy

For classifiers trained on images of natural scenes (118 images, each serving as one classification category), we measured their accuracy using cross-validated F1 score,

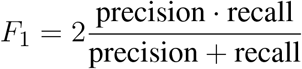

where precision 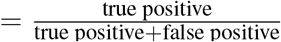 and recall 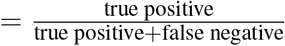. This was computed for each class using `sklearn.metrics.precision recall fscore support` in python. The F1 score ranges from 0 to 1, with higher values indicating better classification and 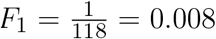 representing chance level.

For classifiers trained on the location of Gabor patches, we incorporated the geometric nature of the task and instead used the distribution of Euclidean distances between the true location and the prediction location of each Gabor patch,

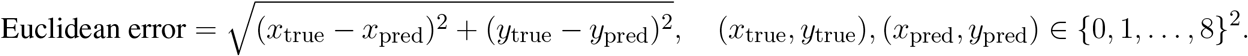

This metric distinguishes between slight misclassifications, where the predicted location is close to the true one, and large misclassifications where the predicted location is many cells away from the true one. Given that the patches were presented at either cell of a 9x9 grid, each Euclidean distance can range from 0 to 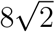, with a chance level of approximately 4.7.

### Alternative classifiers

Alternative machine learning classifiers were implemented using `scikit-learn` in python with the following parameters. Random forest: 10 estimators; SVM: RBF kernel, *C* = 0.5; artificial neural network: multi-layer perceptron with one hidden layer, 10 hidden units, and ReLU activation.

### Computing

All the computations reported in this study were performed on a Lenovo P620 workstation with AMD 3970X 32-Core processor, Nvidia GeForce RTX 2080 GPU, and 512GB of RAM.

## CONFLICT OF INTEREST STATEMENT

The authors declare that the research was conducted in the absence of any commercial or financial relationships that could be construed as a potential conflict of interest.

## AUTHOR CONTRIBUTIONS

EN conceived of the study and supervised the research; TS performed the research; MI conceived of the computational framework to be used an oversaw the design and implementation of NeuroPixelHD; ZZ assisted in the design and implementation of NeuroPixelHD and conducted its theoretical memory analysis; All authors contributed in writing the manuscript.

## FUNDING

The research conducted in this study was partially supported by NSF Award #2239654 to E.N. and DARPA Young Faculty Award, National Science Foundation #2127780 and #2312517, Semiconductor Research Corporation (SRC), Office of Naval Research, grants #N00014-21-1-2225 and #N00014-22-1-2067, the Air Force Office of Scientific Research under award #FA9550-22-1-0253, and generous gifts from Xilinx and Cisco to M.I.

## ACKNOWLEDGMENTS

The authors are thankful to Ali Zakeri and Calvin Yeung for constructive discussions and feedback on the design of the NeuroPixelHD model.

## DATA AVAILABILITY STATEMENT

The datasets analyzed for this study can be found in the Allen Institute Visual Coding - Neuropixels dataset, available at https://portal.brain-map.org/explore/circuits/visual-coding-neuropixels.

## Notes

https://portal.brain-map.org/explore/circuits/visual-coding-neuropixels

### Competing Interest Statement

The authors have declared no competing interest.

### Summary of Updates

Added Supplemental Material

