## supplementary file for "Optimal Decoding of Neural Dynamics Occurs at Mesoscale Spatial and Temporal Resolutions"

### Supplementary Material

#### 1 SUPPLEMENTARY NOTE 1: THEORETICAL ANALYSES

##### 1.1 NeuroPixelHD Memory Analysis

In this section, we analyze the expressive power of HDC and its ability for performing averaging-free computation through a rigorous mathematical analysis of the memory capacity of the NeuroPixelHD classifier.

**Setup.** To simplify the theoretical analyses, we use dot product as the similarity metric, i.e., for any two hypervectors (HVs) $\vec{Q}$ and $\vec{C}_l$, $\delta(\vec{Q}, \vec{C}_l) = \vec{Q}^T \vec{C}_l$. Normalizing this dot product by $\|\vec{Q}\| \cdot \|\vec{C}_l\|$ gives the cosine similarity used in NeuroPixelHD. Recall from the main text that for each trial $k$, the trial HV is computed as $\vec{V}^k = \sum_{t \in \mathcal{T}} \vec{K}^k(t)$ where $\mathcal{T} = \{1, \ldots, M\}$ is the set of all time bins,

$$\vec{K}^k(t) = \Big[ \sum_{\substack{i \in \mathcal{P} \\ n_i^k(t) \neq 0}} n_i^k(t) \vec{S}_i \otimes \vec{H}_+ + \sum_{\substack{i \in \mathcal{P} \\ n_i^k(t) = 0}} \vec{S}_i \otimes \vec{H}_- \Big] \otimes \vec{T}^k(t),$$

$\mathcal{P} = \{1, \ldots, N\}$ denotes the set of neurons, $n_i^k(t)$ is the number of spikes for neuron $i$ in trial $k$ and time bin $t$, $\vec{S}_i$ is the spatial HV for neuron $i$, $\vec{T}^k(t)$ denotes the time HV of time bin $t$ of trial $k$, $H_+$ is the HV indicating spiking activity, and $H_-$ is the HV indicating the absence of spikes. We explicitly parameterize the time HV by $k$ to emphasize that they are generated independently across trials. Then, to classify trials into behavioral categories, we construct a class HV $\vec{C}_l$ for each class $l$ by bundling all trial HVs belonging to that class: $\vec{C}_l = \sum_{k \in \mathcal{C}_l} \vec{V}^k$, where $\mathcal{C}_l$ denotes the set of trial indices belonging to class $l$. This method of training allows for a more concise analysis and is a simplification of the adaptive approach we use in NeuroPixelHD, where the latter leads to a weighted version of the former with negative weights depending on misclassifications during adaptive training.

**Memorization and decoding.** To decode the neural activity HV $\vec{K}^k(t)$ back to its constituent set of neural spike counts $\{n_i^k(t)\}_i$, we may retrieve each $n_i^k(t)$ via

$$\delta(\vec{K}^k(t), \vec{S}_i \otimes \vec{T}^k(t) \otimes \vec{H}_+) \approx D n_i^k(t) \qquad \text{(S1)}$$

where $D$ is the hyperdimension. This can be shown using the dissociation property of binding. To simplify the analysis, assume that the spatial and temporal HVs are nearly orthogonal among themselves and to each other. Then, since binding preserves dissimilarity, $\delta(\vec{S}_i \otimes \vec{T}^k(t), \vec{S}_{i'} \otimes \vec{T}^{k'}(t')) \approx 0$ for $(i, k, t) \neq (i', k', t')$. Now, consider a case where $n_i^k(t) \neq 0$. Dropping the fixed $k$ and $t$ indices to simplify notation, we have

$$\begin{aligned}
\delta(\vec{K}, \vec{S}_i \otimes \vec{T} \otimes \vec{H}_+) &= \sum_{\substack{j \in \mathcal{P} \\ n_j \neq 0}} \delta(n_j \vec{S}_j \otimes \vec{H}_+ \otimes \vec{T}, \vec{S}_i \otimes \vec{H}_+ \otimes \vec{T}) + \sum_{\substack{j \in \mathcal{P} \\ n_j = 0}} \delta(\vec{S}_j \otimes \vec{H}_- \otimes \vec{T}, \vec{S}_i \otimes \vec{H}_+ \otimes \vec{T}) \\
&\overset{(a)}{\simeq} D n_i + \sum_{\substack{j \in \mathcal{P} \\ j \neq i}} \max\{1, n_j\} \mathcal{N}(0, D) \\
&= D n_i + \mathcal{N}(0, (\|\vec{n}\|_2^2 - n_i) D)
\end{aligned}$$

$$\simeq Dn_i + \mathcal{N}(0, \|\vec{n}\|_2^2 D)$$
$$= D\big[n_i + \mathcal{N}(0, \|\vec{n}\|_2^2/D)\big] \tag{S2}$$

where $\mathcal{N}(\mu, \sigma^2)$ denotes a normally distributed random variable with mean $\mu$ and variance $\sigma^2$, and $\vec{n}^k(t) = [\max\{1, n_1^k(t)\}\ \max\{1, n_2^k(t)\}\ \cdots\ \max\{1, n_N^k(t)\}]^T$. The approximate equality $(a)$ results from the fact that $\delta(n_i \times \vec{S}_i \otimes \vec{T} \otimes \vec{H}_+, \vec{S}_i \otimes \vec{T} \otimes \vec{H}_+) = n_i D$ and other terms in the similarity between two nearly orthogonal HVs each contribute a noise term that can be approximated by a normally distributed random variable with mean 0 and variance $D$, scaled by the magnitude of the neural activities ($n_j$ for $n_j > 0$, and 1 for $n_j = 0$). The case where $n_i = 0$ is similar, except that only noise is produced: $\delta(\vec{K}, \vec{S}_i \otimes \vec{T} \otimes \vec{H}_+) \simeq N(0, (\|\vec{n}\|_2^2 D))$. Analogous computation of the similarity can be done with trial HV $\vec{M}^k$ and each class HV $\vec{C}_l$. In particular:

$$\delta(\vec{M}^k, \vec{S}_i \otimes \vec{T}^k(t) \otimes \vec{H}_+) \simeq D\Big[n_i^k(t) + \mathcal{N}\big(0, \sum_{t \in \mathcal{T}} \|\vec{n}^k(t)\|_2^2/D\big)\Big], \tag{S3}$$

and, for $i \in \mathcal{C}_l$,

$$\delta(\vec{C}_l, \vec{S}_i \otimes \vec{T}^k(t) \otimes \vec{H}_+) \simeq D\Big[n_i^k(t) + \mathcal{N}\big(0, \sum_{t \in \mathcal{T}, k \in \mathcal{C}_l} \|\vec{n}^k(t)\|_2^2/D\big)\Big]. \tag{S4}$$

Because of the discreteness of neural activity $n_i^k(t)$ as spike counts, we round the similarity as the final retrieved signal:

$$\hat{n}_{i,t}^{(k)} = \lfloor \delta(\vec{C}_l, \vec{S}_i \otimes \vec{T}^k(t) \otimes \vec{H}_+)/D \rceil. \tag{S5}$$

As noise accumulates with each bundling operation, performing a complete decoding directly from a class HV to each neural activity $n_i^k(t)$ would require a large hyperdimension $D$ to ensure that the signal-to-noise ratio is sufficiently high for producing accurate results. To avoid this issue, one may use intermediate codebooks, so that the decoding can be hierarchical, effectively adding noise reduction during the decoding process. A similar alternative is via iterative noise cancellation method introduced in Poduval et al. (2022): while retrieving multiple entries of $n_i^k(t)$ in parallel, one can improve the prediction of each entry by subtracting the HV introduced by the other entries from the composite HV to effectively reduce the noise they introduce. This process can be iteratively performed to all entries to improve the quality of their decoding.

**Memory accuracy.** To measure the memory accuracy and capacity of a class HV $C_l$, we evaluate its ability to accurately recall neural activity $n_i^k(t)$. Note that this differs from "prediction accuracy", where many class HVs are compared with a query HV to predict whether or not it belongs to a class. As we use rounding at the end of our retrieval, the accuracy of the class HV $C_l$ for each term $n_i^k(t), i \in \mathcal{C}_l$ can be approximated by

$$Acc_i^k(t) = \Pr\left\{\hat{n}_i^k(t) = n_i^k(t)\right\}$$
$$= 1 - 2\Phi\Big(-\frac{1}{2\sqrt{L}}\Big)$$

where $\Phi$ denotes the cumulative distribution function of the standard normal distribution and $L = \sum_{t\in\mathcal{T},k\in\mathcal{C}_l} \|\vec{n}^k(t)\|_2^2/D$ is the variance derived in Eq. S4. Notice that the accuracy for a certain term depends on the norm of all terms involved. Therefore, the average accuracy across all terms is the same: $\mathrm{Acc} = 1 - 2\Phi(-1/(2\sqrt{L}))$. Finally, note that this is a rough estimation of the accuracy from direct recalls while the optimizations mentioned previously may improve the total accuracy.

**Memory capacity.** Following the methodology in (Frady et al., 2018), we define the memory capacity of NeuroPixelHD as its information content: the mutual information between true inputs (spike counts) and those retrievable from each model (class HV) $\vec{C}_l$. For any fixed $(i, k, t)$, with an abuse of notation, Let $n$ be a (continuous) Gaussian approximation of the actual neural spike count and $\hat{n}$ be the Gaussian approximation of the corresponding predicted output from the model in (S5). This discrete to continuous approximation can be quite accurate if, e.g., each discrete spike count follows a Poisson distribution with a rate $\lambda \gtrsim 5$. The mutual information between $n$ and $\hat{n}$ can then be computed via $I_n = D_{KL}(p(\hat{n}, n)\|p(\hat{n})p(n))$ where $D_{KL}$ denotes the KL divergence. Assuming that the joint distribution $p(\hat{n}, n)$ is also Gaussian, the correlation $\rho$ between $p(\hat{n})$ and $p(n)$ would be sufficient to compute $I_n$, as $I_n = -\frac{1}{2}\log_2(1-\rho^2)$ (Gel'fand and Yaglom, 1957). For NeuroPixelHD, correlation coefficient $\rho$ can be derived as $\rho = \frac{n^2}{\sqrt{n^2(n^2+L)}} = \frac{n}{\sqrt{(n^2+L)}}$ (Borga, 2001). Thus, the total information content of each class HV $\mathcal{C}_l$ is given by

$$I = \frac{1}{2} \sum_{k\in\mathcal{C}_l,t\in\mathcal{T},i\in\mathcal{P}} \log_2\left(\left[n_i^k(t)\right]^2/L + 1\right).$$

### 2 SUPPLEMENTARY TABLES AND FIGURES

#### 2.1 Figures

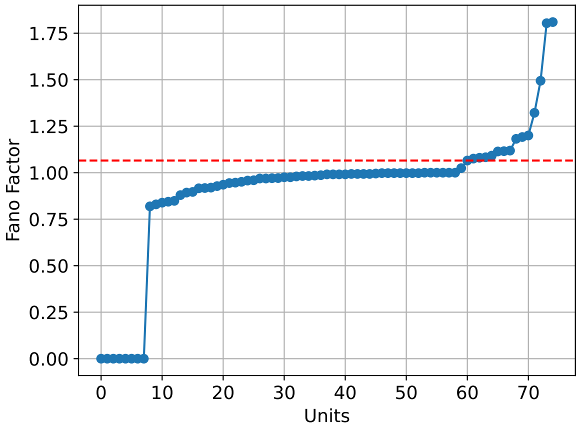


(1a)

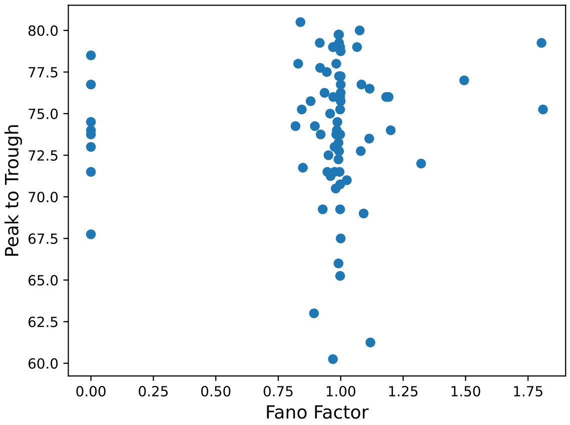


(1b)

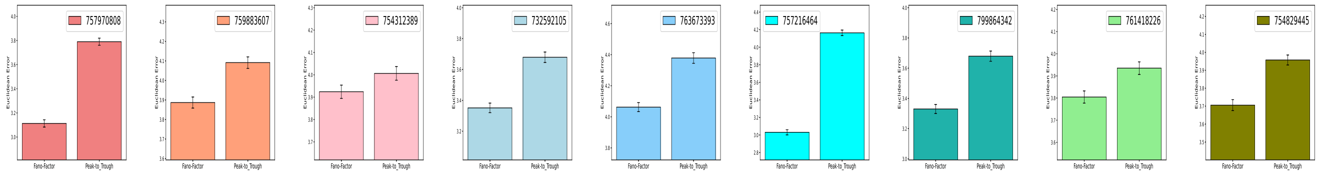


(1c)

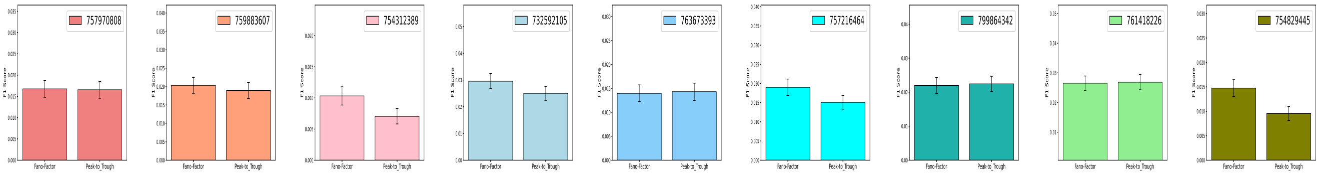


(1d)

Figure S1: **Clustering neurons into putatively excitatory and inhibitory populations based on spike width and Fano factor lead to significantly different outcomes. (1a)** Sorted Fano factors for neurons in a sample brain region (V1) for a randomly selected mouse. The red line represents the threshold (80th percentile) used to distinguish between fast spiking (larger Fano factor, putatively inhibitory) and regular spiking (smaller Fano factor, putatively excitatory) neurons. **(1b)** Scatter plot depicting the absence of any apparent relationship between the Fano Factor and Peak-to-Trough metrics for neurons in the VISp region. **(1c)** Euclidean errors of NeuroPixelHD classification of Gabor locations at the population level spatial resolution. The left and right bars in each panel correspond to population classification based on Fano factor and duration of spike peak to trough, respectively. Error bars represent 1 s.e.m. **(1d)** Similar to (1c) but for F1 scores of classifying images of natural scenes.

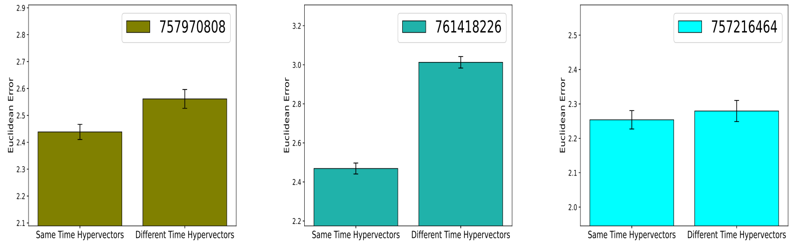


(2a)

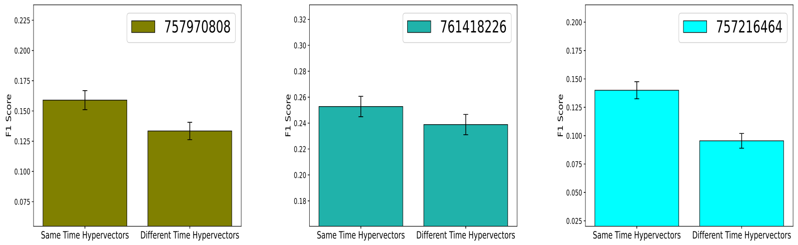


(2b)

Figure S2: **Comparing the results of NeuroPixelHD when using the same set of time HVs for all trials versus defining a unique set of time HVs for each trial. (2a)** Euclidean errors of NeuroPixelHD classification for Gabor locations at the neuron level spatial resolution and 125ms temporal resolution for three randomly selected mice. In each panel, the left and right bars represent the classification outcomes achieved by using a consistent time HV for all trials and using distinct time HVs for individual trials, respectively. **(2b)** Similar to (2a) but for F1 scores of classifying images of natural scenes.
